# A Jacob/nsmf gene knockout does not protect against hypoxia- and NMDA-induced neuronal cell death

**DOI:** 10.1101/2022.12.20.521175

**Authors:** Guilherme M. Gomes, Julia Bär, Anna Karpova, Michael R. Kreutz

## Abstract

Jacob is a synapto-nuclear messenger protein that encodes and transduces the origin of synaptic and extrasynaptic NMDA receptor signals to the nucleus. The protein assembles a signalosome that differs in case of synaptic or extrasynaptic NMDAR activation. Following nuclear import Jacob docks these signalosomes to the transcription factor CREB. We have recently shown that amyloid-beta and extrasynaptic NMDAR activation triggers the translocation of a Jacob signalosome that results in inactivation of the transcription factor CREB, a phenomenon termed Jacob-induced CREB shut-off (JaCS). JaCS contributes to early Alzheimer’s disease pathology and a gene knockout of nsmf, the gene encoding Jacob, protects against amyloid pathology. Given that extrasynaptic activity is also involved in acute excitotoxicity, like in stroke, we asked whether nsmf gene knockout will also protect against acute insults, like oxygen and glucose deprivation and excitotoxic NMDA stimulation. Here we show that organotypic hippocampal slices from wild-type and nsmf^-/-^ mice display similar degrees of degeneration when exposed to either oxygen glucose deprivation or 50 µM NMDA incubation. This lack of neuroprotection indicates that JaCS is mainly relevant in conditions of low level chronic extrasynaptic NMDAR activation that results in cellular degeneration induced by alterations in gene transcription.

## MAIN TEXT

Disruption of cAMP-responsive element-binding protein (CREB) transcriptional activity, a master regulator of cell survival and plasticity-related gene expression, is a hallmark of neurodegeneration^1^. Long-lasting dephosphorylation of CREB at serine 133, termed CREB shut-off, results in early synaptic dysfunction, contributes to pathology and eventually neuronal cell death. It is elicited by sustained activation of extrasynaptic N-methyl-D-aspartate-receptors (NMDAR). Glutamate spillover to peri- and extrasynaptic sites causes in conjunction with binding of amyloid-β (Aβ) detrimental activation of extrasynaptic NMDAR at early stages of Alzheimer’s disease (AD). In previous work we found that the messenger protein Jacob encodes and transduces the synaptic or extrasynaptic origin of NMDAR signals to the nucleus^2^. In response to cell survival and plasticity-related synaptic NMDAR stimulation, macromolecular transport of Jacob from synapses to the nucleus docks the extracellular signaling-regulated kinase (ERK) to the CREB complex which results in sustained CREB phosphorylation at serine 133^2^. Following disease-related activation of extrasynaptic NMDARs, Jacob associates with protein phosphatase-1γ (PP1γ) and induces dephosphorylation and transcriptional inactivation of CREB (Jacob-induced CREB shut-off (JaCS)^3^). Binding of the adaptor protein LIM domain only 4 (LMO4) distinguishes extrasynaptic from synaptic NMDAR signaling and determines the affinity for the association with PP1γ^3^. This mechanism contributes to transcriptional inactivation of CREB in the context of early synaptic dysfunction in AD^3^. Accordingly, Jacob protein knockdown attenuates Aβ-induced CREB shut-off induced via activation of extrasynaptic NMDARs and nsmf gene knockout is neuroprotective in a transgenic mouse model of AD^3^. Collectively the data suggest that long-distance protein transport from extrasynaptic NMDAR to the nucleus is part of early AD pathology and that Jacob docks a signalosome to CREB that is instrumental for CREB shut-off.

We now asked whether this mechanism is also relevant in cell death induced by acute excitotoxic insults, like those resulting from traumatic brain injury and stroke^4^. While the molecular underpinnings that drive cell death might differ between acute and chronic neurodegenerative insults, activation of extrasynaptic NMDAR appears to be fundamental in both conditions. To tackle this question, we employed organotypic hippocampal slice cultures (OHSC) of wild-type (wt) and nsmf knockout mice^5^, and submitted them to two well-established protocols to study stroke-like excitotoxic insults. We predicted that the nsmf gene knockout would have a neuroprotective effect on OHSC exposed to either oxygen and glucose deprivation (OGD) or bath application of high doses of NMDA^6.^

In the first set of experiments, OHSC from wt and nsmf^-/-^ mice were submitted to OGD for 30 min and cell death was assessed via monitoring propidium iodide (PI) uptake at different intervals after the insult (3 h, 8 h, 12 h and 24 h; for detailed methods see Additional File 1). PI is a red-fluorescent nuclear counterstain not permeant to living cells, thus the increase in fluorescence provides a read out of cell death. Statistical analysis revealed that exposure of OHSC to 30 min OGD induces strong cell death in the CA1 and CA3 subregions of the hippocampus of wt slices, as early as 3 h after the insult (Figure 1A, C, D, E, Two-way repeated measures ANOVA, time x OGD CA1 F_(12,140)_=25.43, p<0.0001; CA3 F_(12,140)_=10, p<0.0001; DG F_(12,140)_=11.42, p<0.0001). Cell death was also detected in the dentate gyrus (DG), although to a lower degree when compared to the other subregions, which is in line with several studies applying OGD^7,8^. Surprisingly, regardless of the subregion analyzed, no difference in cell death between wild-type and nsmf^-/-^ slices was observed at all time points and in all subregions analyzed (Figure 1A, C, D, E, Mixed-effects model analysis CA1 nsmf^+/+^ x nsmf^-/-^ F_(1,19)_=0.003, p=0.9057; CA3 F_(1,19)_=0.3232, p=0.5763; DG F_(1,19)_=0.1593, p=0.6942).

**Fig 1.**
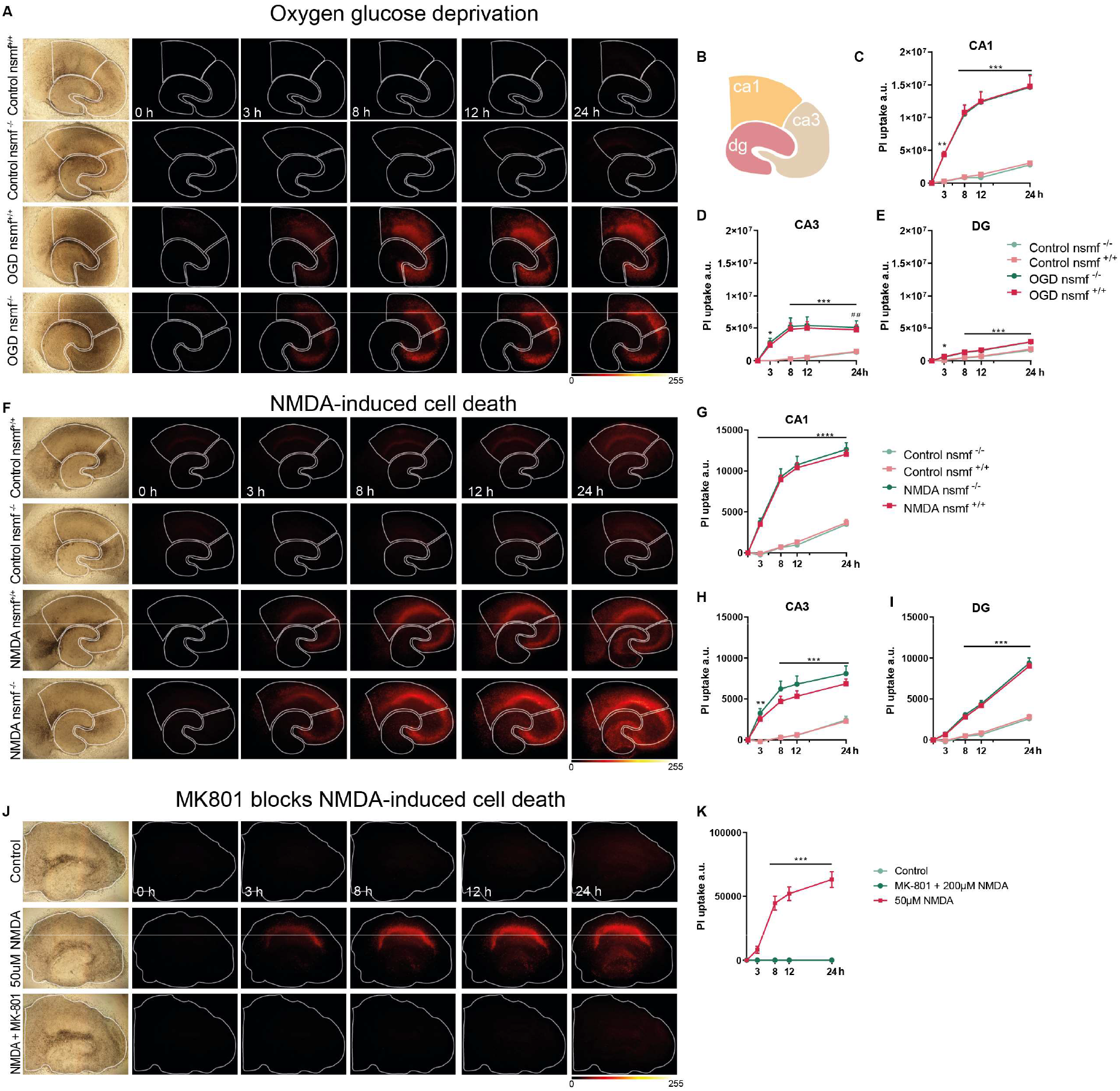
Jacob/nsmf gene knockout does not protect against OGD- and NMDA-induced cell death. **(A-E)** Oxygen glucose deprivation (OGD) induces cell death in organotypic hippocampal slice culture irrespective of mice genotype. **(A)** Bright field and fluorescent images of propidium iodide (PI) signal in organotypic hippocampal slices from wild-type (nsmf^+/+^) and Jacob/nsmf constitutive knock-out animals (snmf^-/-^) 0, 3, 8, 12, and 24 h after 30 min OGD and control. **(B)** Scheme representing CA1, CA3, and DG areas. **(C-E)** Graphs representing the degree of PI uptake in arbitrary units (A.U.) over time (h) after OGD insult. The OGD protocol induced cell death to the same degree in CA1(C), CA3 (D), and DG (E) irrespective of genotype. N= Control nsmf^+/+^:11; OGD nsmf^+/+^:13; Control nsmf^-/-^:7; OGD nsmf^-/-^:8 slices per group. ** p<0.01, *** p<0.001 OGD Jac^+/+^ x control Jac^+/+, ##^ p<0.01 OGD Jac^-/-^ x control Jac^-/-^ by repeated measures (RM) two-way ANOVA followed by Bonferroni’s multiple comparisons test. Data represented as mean ± SEM. **(F-I)** Acute NMDA (50 µM) treatment induces cell death in OHSC irrespective of genotype. **(F)** Brightfield and PI signal in organotypic hippocampal slices from Jac^+/+^ Jac^-/-^ animals after 0, 3, 8, 12, and 24 h post treatment with 50 µM NMDA or control. **(G, H, I)** Graphs representing the degree of PI uptake (A.U.) over time (h) after treatment with 50 µM NMDA. 50 µM NMDA induced cell death to the same degree in CA1, CA3, and DG irrespective of genotype. N= Control nsmf^+/+^:8; NMDA nsmf^+/+^:15; Control nsmf^-/-^:6; NMDA nsmf^-/-^:10 slices per group. ** p<0.01, *** p<0.001, **** p<0.0001 NMDA nsmf^+/+^ x control nsmf^+/+^ by RM two-way ANOVA followed by Bonferroni’s multiple comparisons test. Data represented as mean ± SEM. **(J-K)** MK-801 blocks NMDA-induced cell death in OHSC from C57BL6/J mice. **(J)** Brightfield and PI signal from OHSC 0, 3, 8, 12 and 24 h after treatment with 50 µM NMDA, co-application of MK-801 + 200 µM NMDA, or control. (K) The co-application of MK-801 completely abolished the effects of NMDA treatment. N=Control:4; NMDA:8; NMDA+MK-801:4 slices per group. *** p<0.001 vs. control by RM two-way ANOVA followed by Bonferroni’s multiple comparisons test. Data represented as mean ± SEM. Lookup table indicates the pixel intensities from 0 to 255.

In the next set of experiments, we assessed whether nsmf gene knockout confers protective effects on OHSC incubated with 50 µM NMDA. Statistical analysis revealed that 30 min bath application of NMDA induces strong cell death in wild-type slices over time, as indicated by the increase in PI uptake in all subregions as early as 3 h (Figure 1F, G, H, I, Two-way repeated measures ANOVA, time x OGD CA1 F_(12,140)_=47.07, p<0.0001; CA3 F_(12,140)_=15.68, p<0.0001; DG F_(12,132)_=54.57, p<0.0001). NMDA-induced cell death reached a plateau in both CA1 and CA3 subregions 8 h after NMDA bath application (Fig 1G, H), while cell death reached its peak at 24 h in DG during the examined time period (Fig 1I). Similar to the OGD experiments, no difference in cell death between wild-type and nsmf^-/-^ slices was observed at all time points and in all subregions analyzed (Figure 1F, G, H, I, Mixed-effects model analysis CA1 nsmf^+/+^ x nsmf^-/-^ F_(1,23)_=1.097, p=0.3058; CA3 F_(1,23)_=1.602, p=0.2182; DG F_(1,23)_=0.2236, p=0.6408). Lastly, as a control experiment, we co-applied the NMDAR antagonist MK-801 to OHSC in order to confirm that NMDA-induced cell death occurs via activation of NMDA receptors. Statistical analysis revealed that co-application of MK-801 with NMDA completely abolished PI uptake, at all time-points (Figure 1J, K, Two-way repeated measures ANOVA groups x time F_(1,6)_=0.3347, p=0.5840).

Here we showed that both OGD and NMDA protocols induce cell death in wild-type and nsmf^-/-^ OHSC slices to the same extent, suggesting that JaCS is not involved in acute excitotoxic insults. Excessive entry of Ca^2+^ via NMDARs causes disruption of mitochondrial calcium homeostasis, leading to neuronal cell death by apoptosis^9^. In the face of an acute excitotoxic insult, production of reactive oxygen species and breakdown of the mitochondrial membrane potential are the probable culprits for neurodegeneration^6^. In conclusion, JaCS appear to be relevant in scenarios where activation of extrasynaptic NMDARs builds up slowly, is chronic and results in cellular degeneration due to alterations in gene transcription.

## Abbreviations

NMDAR: N-methyl-D-aspartate receptor
AD: Alzheimer’s Disease
ERK: extracellular signaling-regulated kinase
nsmf: NMDAR synaptonuclear signaling and neuronal migration factor
PP1γ: protein phosphatase-1γ
LMO4: LIM domain only 4
Aβ: amyloid-beta peptide.
OHSC: organotypic hippocampal slice cultures
WT: Wild-type
CREB: cAMP response element-binding protein
NMDA: N-methyl-D-aspartate
OGD: oxygen and glucose deprivation
DG: Dentate Gyrus
CA1: Cornus ammonis 1
CA3: Cornus ammonis 3
ANOVA: Analysis of variance
PI: Propidium Iodide

---

**Additional file 1**. Extended materials and methods, detailed information on statistics.

## Acknowledgments

We would like to thank Corinna Borutzki, Stefanie Hochmuth and Monika Marunde for excellent technical assistance. We also thank Dr. Klaus Reymann for provision of the fluorescence microscope and for help with establishing the protocols used in this study.

## Authors’ Contributions

GMG and JB designed and carried out all the experiments, performed statistical analysis, and wrote the manuscript. AK designed and carried out experiments. MRK conceived and supervised experiments, wrote the manuscript and accrued funding. All authors read and approved the final manuscript.

## Availability of data and materials

Please contact the corresponding author for data requests.

## Declarations

## Ethics approval and consent to participate

Experiments were conducted following ethical animal research standards defined by the German Law/European directive and approved by the Landesverwaltungsamt Saxony-Anhalt (Referat 203, Verbraucherschutz und Veterinärangelegenheiten). The competent authority follows the advice of an official animal welfare committee of the Federal State of Saxony-Anhalt, Germany.

## Consent for publication

Not applicable.

## Additional File 1

### Extended materials and methods, detailed information of statistics Animals

Male C57BL/6J mice and Jacob/nsmf knockout mice^1^ were bred and maintained in the animal facility of the Leibniz Institute for Neurobiology, Magdeburg, Germany. Animals were housed in groups of up to 5 in individually ventilated cages (IVCs; Green line system, Tecniplast) under controlled environmental conditions (22 °C +/-2 °C, 55 % +/-10 % humidity, 12 h light-dark cycle, lights on at 06:00 am). Food and water were provided *ad libitum*.

### Culturing of murine hippocampal slices (OHSC), stimulation

OHSC were prepared according to a previously published protocol^2^. Slices were prepared from P7-P9 Jacob/Nsmf knockout mice and wild-type littermates from heterozygous breeding and cultured for approximately 11 day. OHSC culturing medium consisted of 50% minimal essential medium (Gibco), 25% heat inactivated horse serum (Gibco), 25mM glucose, 2mM glutamine, 25 mM HEPES, 1x B27 and 1x pen/strep, buffered in HBSS+/+ (containing bivalent cations). For experiments with MK-801 slices were prepared from P12 C57BL6/J mice. Before cell death experiments, the viability of slices was assessed by adding 2 μM propidium iodide (PI) for several hours directly into the culture medium and only slices without (or very little, <10 cells) PI fluorescence were used for subsequent experiments.

For OGD-induced cell death experiments, membranes with OHSC were transferred to 1 ml OGD medium (Ringer solution [124 mM NaCl, 4.83 mM KCl, 1.3 MgSO_4_, 1.97 mM CaCl_2_, 1.21 mM KH_2_PO_4_, 25.6 mM NaHCO_3_ pH 7.4] with 10 mM mannitol or 10 mM glucose as a control) with 3 μM PI. Next, cultures were exposed to 10 min of 95 % N_2_/5% CO_2_ gas flow in a hypoxic chamber (Billupsand, Rothenberg) and kept in the reduced O_2_ conditions for 30 min at 35 °C. Afterwards, the membranes were moved back to their previous medium containing 3 μM PI and imaged at the indicated time points.

For NMDA-induced cell death experiments, OHSC were transferred to the medium containing 50 μM NMDA (Sigma-Aldrich) and 3 μM PI for 30 min. Afterwards, membranes were moved back to their previous medium containing 3 μM PI and imaged at the time points indicated in the figure legend. In order to assure that the observed cell death was NMDAR-dependent, slices were pre-incubated with 10 μM MK-801 (Sigma-Aldrich) for 1 h and during the 200 µM NMDA exposure.

To assess the cell death, the images of PI fluorescence, as well as brightfield images, were acquired on a Zeiss Axioskop 2 with the 4x objective using Nikon NIS software. Analysis was performed in Fiji^3^. The CA1, C3, and DG regions of interest (ROIs) were defined manually based on the brightfield image. Mean pixel intensity was measured in grey-scale images of the PI channel, and individually baseline-corrected for time point 0 (directly after the experiment). The analysis was done by an experimenter blind to genotype and treatment group. For representative images, PI images of different time points were aligned using the Fiji plugin linear stack alignment with SIFT and default settings in the rigid mode. Represented images are contrast-enhanced (equally for all groups).

### Data Analysis

The results are presented as mean +/-SEM. Data were analyzed using two-way analysis of variance (ANOVA) followed by Bonferroni post-hoc test, and a mixed-effect model analysis. Statistical significance was considered as p<0.05. Data was analyzed and plotted using GraphPad Prism v9.0.2 (Graph Pad Software, San Diego, USA).

